# Shamonda virus EU1 in Northern Italy, August 2026

**DOI:** 10.64898/2026.09.17.752072

**Authors:** Letizia Ceglie, Alexander Tavella, Debora Dellamaria, Ilaria Belfanti, Erika Rampazzo, Florian Beikircher, Gelindo Berasi, Mattia Fustini, Christoph Gasser, Judith Kristler Pallhuber, Julia Poernbacher, Thomas Schwienbacher, Elisa Palumbo, Alessandro Sartori, Luca Tassoni, Isabella Monne, Giovanni Cattoli, Paola De Benedictis

## Abstract

**Background:** Following the emergence of a novel Shamonda-like virus (SHAV) in Europe, suspected clinical cases in dairy cattle were reported in Northeastern Italy. Investigation was required in dairy cattle in Northeastern Italy.

**Methods:** Samples were collected from 28 farms displaying fever and reduced milk yield and were screened using molecular methods (PanSimbu real-time RT-PCR followed by Sanger sequencing and differential PCRs for SBV/BTV). Positive samples were subsequently confirmed by Sanger sequencing and selected samples underwent whole-genome sequencing using a metagenomic approach.

**Results:** Twenty-four among sera and blood samples from 12 farms tested positive for SHAV. WGS confirmed that Italian strains share maximum identity with the European SHAV-EU1 clade.

**Limitations:** This study concerns the results of clinical investigations on suspected cases in the two autonomous provinces of Trento and Bolzano. At present, it is not possible to infer on the circulation of the virus in other Italian provinces.

**Conclusions:** This study documents the first detection and genetic characterization of SHAV-EU1 in Italian cattle, indicating active circulation of this virus across Europe. Its identification in two north-eastern Italian provinces does not rule out wider, undetected circulation. In this regard, assessing its geographical extent requires careful evaluation, just as much remains to be discovered about its host range and the severity of symptoms.

## Introduction

Shamonda virus (SHAV) is a single-stranded, negative-sense RNA virus belonging to the family *Peribunyaviridae*, order *Bunyavirales*, and the Simbu serogroup. Its genome is composed of three segments, designated Large (L), Medium (M), and Small (S). First isolated in Nigeria in the 1960s from cattle and *Culicoides* biting midges, SHAV was historically restricted to Africa, Asia, and Australia. However, in summer 2026, a novel “Shamonda-like” virus, hereinafter called European Shamonda virus (SHAV) unexpectedly emerged across Central and Western Europe, including France, Germany, Switzerland, the Netherlands, Austria, Belgium, and Liechtenstein (1,2). Initial clinical investigations in dairy herds revealed an acute, non-fatal syndrome marked by transient fever (>40 °C), severe diarrhoea, lethargy, anorexia, and substantial drops in bulk milk production ranging from 15% to 75% (2–4). Routine diagnostic assays for established arboviruses, such as Schmallenberg virus (SBV) and bluetongue virus (BTV), yielded negative results. The causative pathogen was subsequently identified using pan-Simbu real-time RT-qPCR and untargeted Nanopore metagenomic sequencing, leading to whole-genome recovery and taxonomic classification within *Orthobunyavirus schmallenbergense* (4,5). Transmitted primarily by *Culicoides* midges, there are currently no specific vaccines or treatments available (1,6). Its rapid spread across Europe has led to the identification of two co-circulating clades, designated SHAV-EU1 and SHAV-EU2. The SHAV-EU1 clade has so far been detected in France, Switzerland, Germany, and Austria, and exhibits the highest nucleotide identity with historical SHAV isolates in the L and M segments (4,5). The SHAV-EU2 clade has been found in northern Germany, Belgium, the Netherlands, and northwestern France (1).

The present work describes the first detection in Italian dairy herds of the European SHAV and provides the preliminary molecular characterization and phylogenetic analyses of representative isolates as well as the first evidence of its circulation in Italy.

## Materials and methods

Between late August and early September 2026, overall 48 K3EDTA blood samples, 7 sera and 5 organs from aborted fetuses were collected from 28 dairy farms in Northeastern Italy and sent to IZSVe to investigate the presence of the European SHAV. All collected samples belonged to stabled cattle suffering from fever, diarrhea, decreased milk production and prostration. All farms are located in the two Autonomous Provinces of Bolzano and Trento, at an altitude ranging from 450 to 1,400 m a.s.l..

Viral RNA extraction was semi-automatically performed using the MagMAX CORE Nucleic Acid Purification Kit and the King Fisher**™** 96 Flex purification system (Thermo Fisher Scientific). A universal PanSimbu S-segment-based real-time RT-PCR (7) and the specific commercially available ID Gene™ SHLV Duplex kit (Innovative Diagnostics) were applied to test the extracted RNAs. In parallel, differential diagnostic testing for Schmallenberg virus and bluetongue virus was performed using a S3-Schmallenberg specific real-time RT-PCR and a NS3-based panBTV commercial assay (LSI, Thermo Fisher Scientific) (8). Samples testing positive by PanSimbu realtime RT-PCR (7) were subsequently confirmed by Sanger sequencing. Two positive samples, originating from two different farms located in the provinces of Trento and Bolzano, respectively, were selected for whole-genome sequencing (WGS) using a metagenomic NGS approach. Total RNA was purified and concentrated using the RNeasy MinElute Cleanup kit (Qiagen). Removal of abundant ribosomal RNA (rRNA) and globin RNA, as well as library preparation, were carried out with the Illumina Stranded Total RNA Prep, Ligation with Ribo-Zero Plus according to manufacturer’s instructions (Illumina). Samples were sequenced on an Illumina MiSeq platform using a MiSeq Reagent Kit v3 in 2×301 bp paired-end mode (Illumina). The consensus sequence of the whole genome was obtained using an in-house Nextflow pipeline (1). Sequencing adapters and low quality (<Q20) 3’ ends were trimmed with Cutadapt version 4.5 (2). All reads shorter than 80 bp after the trimming were discarded. Reads quality was assessed with FastQC v.0.12.1 (3). Assembly was performed using a reference-based approach, in which reads were mapped with BWA 0.7.17 (4) and the obtained alignment was corrected and improved using GATK version 4.4.0.0 (5). Variant calling was performed using LoFreq v.2.1.5 (9–13) and the consensus sequence was generated with an in-house Python script. For sample 26/114900, horizontal coverage was 95% and vertical coverage between 50x and 120x. For sample 26/114491, horizontal coverage was 98% and vertical coverage between 80x and 120x. The references used for read assembly were selected from a custom-built NCBI dataset using BLAST 2.15.0. The selected accession numbers were, respectively for the two samples: S=OZ516536.1, M=OZ516535.1, L=OZ516531.1, S=OZ516536.1, M=PZ831159.1, L=OZ516534.1.

The obtained sequences were analysed to reconstruct the phylogenetic relationships between the Italian viruses and other related Orthobunyaviruses. Closely related sequences were identified through BLAST searches in NCBI (last accessed 14 September 2026) (14). All sequences isolated in Germany, France, and Switzerland in 2026 and publicly available at the time of the search were included.

Multiple sequences alignments were obtained using MAFFT v.7 and subsequently manually curated and trimmed to retain only the coding sequences (15). Phylogenetic trees were inferred using the maximum-likelihood algorithm implemented in IQ-TREE V3.0.1 (16). The best-fit substitution model for each segment was selected using ModelFinder, as implemented in IQ-TREE, according to the Bayesian Information Criterion (17). The selected models were GTR+F+I+G4 for segment L, TIM3+F+I+G4 for segment M, and TPM2u+I+R2 for segment S. Branch support values were assessed using 1,000 ultrafast bootstrap replicates (18). Phylogenetic trees were visualised using FigTree v1.4.3 .

## Results

Eighteen out of 55 blood and sera samples tested positive by the universal rRT-PCR. The commercial kit confirmed these results and detected 6 additional positive samples. All organs tested negative with both assays. All samples gave negative results in both the specific S3-SBV and the NS3 panBTV assays (8). Twelve out of the 28 surveyed farms turned out positive for SHAV circulation. Epidemiological and clinical data in dairy cattle collected in the two autonomous provinces are summarized in Table 1.

**Table 1.** List of the infected dairy farms in Northern Italy, August-September 2026.

| Case | Date of sampling | Province | Breeds | N. of lactating cows | Drop in milk production<sup>°</sup> (%) | Duration of the drop in milk production (days) | Cows with fever/depression (%) | Duration of diarrhea (days) | Abortion |
| --- | --- | --- | --- | --- | --- | --- | --- | --- | --- |
| 26/120454 | 23/08/2026 | Trento | HF, BS | 65 | 20 | 14 | 5/30 | 6 | 0 |
| 26/114491 | 25/08/2026 | Bolzano | HF, SIM, J | 27 | 80 | 12 | 70/70 | 6 | 0 |
| 26/115226 | 27/08/2026 | Trento | BS | 110 | 30 | 15 | 10/80 | 4 | 1* |
| 26/114851 | 27/08/2026 | Trento | HF | 400 | 10 | 10 | 4/20 | 3 | 0 |
| 26/114893 | 27/08/2026 | Trento | HF | 90 | 5 | 8 | 2/10 | 3 | 0 |
| 26/114900 | 27/08/2026 | Trento | HF | 130 | 15 | 12 | 5/30 | 4 | 0 |
| 26/118701 | 31/08/2026 | Bolzano | BS, HF | 20 | 50 | 14 | 80/80 | 5 | 0 |
| 26/117612 | 01/09/2026 | Bolzano | GA, SIM | 32 | 40 | 14 | 40/50 | 5 | 0 |
| 26/117639 | 02/09/2026 | Bolzano | SIM | 18 | 85 | 10 | 60/80 | 5 | 0 |
| 26/118704 | 03/09/2026 | Bolzano | HF, SIM | 25 | 50 | 12 | 80/80 | 6 | 0 |
| 26/118250 | 03/09/2026 | Bolzano | HF | 33 | 80 | 10 | 60/50 | 5 | 0 |
| 26/119295 | 04/09/2026 | Bolzano | HF | 15 | 50 | 12 | 30/50 | 5 | 0 |

HF = Holstein Friesian; BS = Brown Swiss; SIM = Simmental; J = Jersey; GA = Grey Alpine.
<sup>°</sup> Average drop in milk production in symptomatic cattle.
\* SHAV infection in the foetal organs has been ruled out.

**Figure 1.**
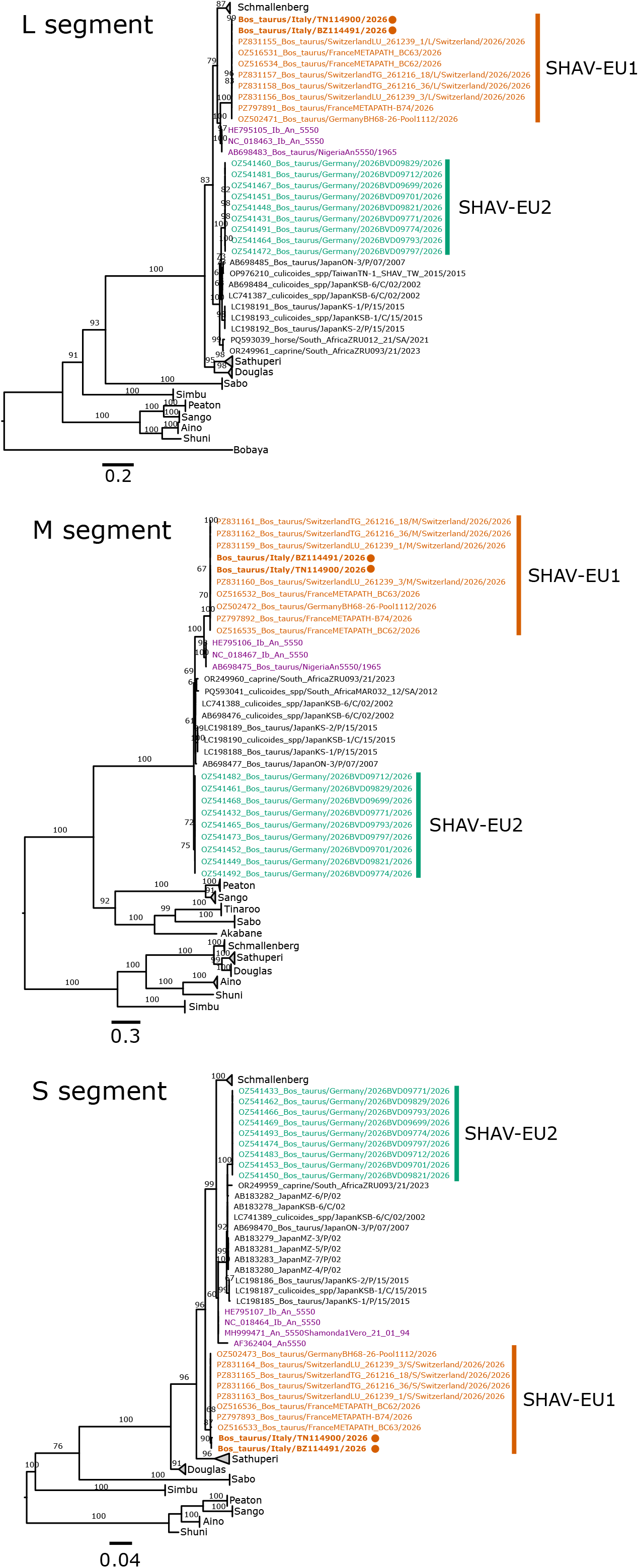
Maximum-likelihood phylogenetic trees of the SHAV sequences and related Orthobunyaviruses (collapsed). Taxa corresponding to the original Ib An 5550 strain are colored in purple. Italian SHAV are marked with a dot

According to the Ct values recorded, 14 of these samples underwent Sanger sequencing that confirmed the presence of Shamonda-like virus. The whole-genome sequences of the Italian SHAV-EU1 viruses have been deposited in GenBank (Accession no. to be assigned). When comparing Italian samples with other European SHAV-EU1 sequences, sequence similarities ranged from 99.9-100% for the L segment, 99.9-100% for the M segment, and 99.6-100% for the S segment. Similarities with the Nigerian SHAV Ib An 5550 reference strain ranged from 92.9- 93.7%, 92.4-92.6%, and 95.6-97.3–XX% for the L, M, and S segments, respectively. Comparing Italian samples with other SHAV, specifically with the clade comprising SHAV-EU2 (collected in Germany in 2026) and Asian and South African SHAV detected between 2002 and 2023 the similarities ranges from 87.5-89.9%, 84.6-87.2%, and 95.9-96.6% for the L, M, and S segments, respectively.

## Discussion

The emergence of unknown pathogens, or pathogens appearing unexpectedly in a specific geographic area or host species, has become a frequent occurrence. Several epizootic outbreaks of orthobunyaviruses have been observed in ruminants in recent decades (8,19), Europe is not exception. Indeed, following the emergence of Schmallenberg virusin 2011 (8), the circulation of at least two lineages of SHAV was reported in Europe in summer 2026 (2,4,5). It is interesting to note that SHAV-EU1 isolates clustered with the Nigerian Shamonda virus (Ib An 5550, isolated from cattle in 1965) in the L and M segments, yet differed in the S segment (20). This makes difficult to determine whether its emergence resulted from a new introduction followed by a high mutation rate or from a reassortment event (4,21,22). Indeed, gene reassortment is a well-known phenomenon for Orthobunyaviruses, and is considered a crucial event underlying viral evolution and the emergence of new strains. Pre-existing population immunity to SBV provides no cross-protection due to M-segment divergence, and concurrent circulation of SHAV variants alongside SBV creates a high potential for genetic reassortment.

The occurrence of epizootic events right on Italy’s borders, combined with reports from veterinarian practitioners regarding lactating cows suffering from fever, diarrhoea, and an unexpected and/or severe drop in milk production, prompted us to adopt a diagnostic approach capable of identifying the emerging SHAV, that was characterised to be SHAV-EU1. This first identification paves the way to investigate several issues that are still unclear regarding this virus.

First, identification in the provinces of Trento and Bolzano alone does not indicate the absence of circulation in more widespread areas of Italy. Structured surveillance that includes the differential diagnosis of SHAV in cases of fever, depression, and a drastic drop in milk production in ruminants is then necessary throughout the country, in order to better understand the disease dynamics and to clearly trace back its history and possibly introduction events.

Second, beyond acute herd morbidity, major veterinary concerns focus on transplacental transmission of Simbu group viruses, which causes abortions, stillbirths, and severe congenital malformations (2,4,23). Although the European SHAV has been associated to abortion in cattle so far, we collected no evidence of these events occurring in the infected farms, nor SHAV has been detected in the organs of the aborted fetuses examined so far. Indeed, the actual impact in cattle’s fertility deserves to be further investigated and quantified, as does the impact on other host species that have hitherto been overlooked. The host range of SHAV observed to date extends to sheep, goats, alpacas, and horses. In these latter, the experience of other European countries highlights the need to include SHAV in the differential diagnosis of neurological syndromes, as viral RNA has been detected in the brain tissue of equine exhibiting neurological signs (24).

Furthermore, the eco-epidemiological aspects of the disease—such as the factors determining the virus’s persistence in nature (e.g., vectors and wildlife)—should not be overlooked, in order to identify risk factors and potential mitigation measures to be implemented.

Finally, although most Simbu serogroup orthobunyaviruses are not currently considered to pose a significant zoonotic risk to humans, the documented or suspected zoonotic potential of related viruses such as Shuni virus and the established ability of Oropouche virus to cause human disease highlight the need for a One Health approach to assess and continuously monitor potential zoonotic risks as epidemiological and ecological conditions evolve (23).

